# Immune responses to human pathogens exposed to simulated Mars conditions

**DOI:** 10.1101/2025.02.05.636683

**Authors:** Tommaso Zaccaria, Özlem Bulut, Anaisa V. Ferreira, Margo Dona, Jeroen D. Langereis, Rob J. Mesman, Joppe Wesseling, Laura van Niftrik, Mihai G. Netea, Petra Rettberg, Kristina Beblo-Vranesevic, Marien I. de Jonge, Jorge Domínguez-Andrés

## Abstract

The identification of health risks associated with long-term crewed missions to Mars is critical for mission planning and crew safety. Human-associated pathogens can be part of the microbiome and are likely to be transported during these missions. This study examines the immunological responses of human immune cells stimulated with non-fastidious bacterial species that cause opportunistic infections, i.e. *Klebsiella pneumoniae* and *Serratia marcescens*, after exposure to simulated Martian conditions, including UV radiation, desiccation and atmospheric pressure. We observed that exposure of the bacteria to these conditions altered cytokine secretion, reactive oxygen species (ROS) production, and phagocytic activity in human peripheral blood mononuclear cells (PBMCs). Specifically, exposure to desiccation reduced cytokines and ROS production, indicating impaired innate immune recognition and stimulation. Notably, the altered immune response was partially restored when desiccated bacteria were regrown in standard media. Flow cytometry revealed decreased bacterial size and complexity of both species post-exposure. These findings indicate that Martian conditions induce bacterial morphological and physiological changes, which could impair immune recognition and response. Expanding these studies to *in vivo* models and a broader range of potentially pathogenic microorganisms is essential to estimate infection risks during Mars missions, which is vital for developing strategies to mitigate infection risks and maintain astronaut health during long-term space travel.

**Importance:** Since Yuri Gagarin’s 1961 flight, human space exploration has expanded, unintentionally transporting microorganisms, including pathogens, into space environments. Our previous studies demonstrated that opportunistic pathogens like *Klebsiella pneumoniae* and *Serratia marcescens* can survive simulated Mars conditions. With upcoming Mars missions, it is crucial to understand how such conditions influence these pathogens and their interaction with the human immune system. This research evaluates immune responses to bacteria pathogens exposed to Martian stressors such as UV radiation and desiccation, revealing significant changes of the immune responses to the exposed bacteria. These findings provide essential insights into the health risks that astronauts may face if infected with Mars-adapted pathogens. Understanding these interactions will help to develop preventive strategies and therapeutic measures, ensuring the safety and health of crew members during long-term missions. Ultimately, this work contributes to the broader objective of safe human exploration and colonization of Mars.

## Introduction

Robotic exploration of Mars has driven extensive engineering and scientific research, including plans for future human missions to the red planet. Most space biology research to date has been conducted in low Earth orbits (LEO), particularly on the International Space Station (ISS), and CubeSats (Zea et al., 2021). Research has shown that microorganisms isolated on the ISS primarily originate from human colonizing flora (Checinska et al., 2015), indicating that similar microorganisms would likely be transported during Mars missions.

The evaluation of microorganisms in spacecrafts has been an ongoing focus over the years (Puleo et al., 1973, Cioletti et al., 1991, Ott et al., 2004) and continues routinely on the ISS (Lang et al., 2017). Since a significant percentage of space station microbes originates from the crew microbiome (Checinska et al., 2015), it is expected that human microbiota will similarly be transported to Mars. Interestingly, certain human pathogens become more virulent in space conditions (Koehle et al., 2023) and show increased resistance to antibiotics (Tierney et al., 2022). As observed on the ISS, pathogenic bacteria species are expected to be present in future Lunar and Martian habitats (Mora et al., 2019). Thus, the risk of infections could be heightened by both altered bacterial virulence factors (Gilbert et al., 2022), and changes in the human immune system, such as immunosuppression during spaceflight (Jacob et al., 2023).

Our previous research demonstrated that *Burkholderia cepacia*, *Serratia marcescens*, *Pseudomonas aeruginosa* and *Klebsiella pneumoniae*, bacterial species that survive and proliferate in a broad range of environmental conditions on Earth, can survive Mars-simulated conditions, even when grown in minimal media with a single carbon source (Domínguez-Andrés et al., 2020, Zaccaria et al., 2024). In this study, we investigated the response of human immune cells to *K. pneumoniae* and *S. marcescens* exposed to Mars-like conditions. These species have been selected for their unique survival traits. We aimed to evaluate whether Mars-simulated conditions affect immune recognition and response to these bacteria. We observed changes in cytokine production, reactive oxygen species (ROS) levels, and phagocytic capacity in response to Mars-exposed *K. pneumoniae* and *S. marcescens*. We also identified morphological changes in these bacteria post-exposure, which may impact their interactions with the immune cells. Our findings enhance the understanding of immune responses to Mars-adapted bacteria, informing strategies to manage infections during long-term missions. These insights are essential for developing targeted therapies and prevention strategies, contributing to preserving astronaut health during space missions.

## Results

We investigated responses of human immune cells to *K. pneumoniae* and *S. marcescens* after exposure to Mars-like conditions, including desiccation, UV radiation, and Martian atmosphere. Our previous work demonstrated that *Burkholderia cepacia*, *Serratia marcescens*, *Pseudomonas aeruginosa* and *Klebsiella pneumoniae* survive in minimal media with a single carbon source and endure Mars-simulated conditions (Domínguez-Andrés et al., 2020, Zaccaria et al., 2024). Based on these results and their pathogenic potential, we selected *K. pneumoniae* and *S. marcescens* for further investigation (Abbas et al., 2024, Khanna et al., 2013). UV doses differed for the two bacterial species, based on the increased resistance to UV irradiation of *S. marcescens* compared to *K. pneumoniae*, which we have highlighted in our previous study (Zaccaria et al., 2024).

### Mars-like conditions decrease the capacity of bacteria to trigger innate and adaptive cytokine production

We first quantified innate and adaptive cytokine release from PBMCs stimulated with Mars-exposed bacteria. *K. pneumoniae* exposed to simulated Mars conditions led to significantly reduced secretion of pro-inflammatory cytokines (IL-1β, IL-6, TNFα) and anti-inflammatory cytokines (IL-1RA, IL-10) compared to controls grown in minimal media (Figure 1A). After 7-day stimulation, Mars condition-exposed *K. pneumoniae* also triggered less T-cell derived cytokines, IL-17 and IL-10, compared to controls (Figure 1B). However, IFNγ and IL-22 production remained unchanged across conditions (Figure 1B). All simulated Mars conditions triggered a statistically significant decrease in the release of TNFα and IL-10, while desiccation did not alter IL-1β, IL-6 or IL-1RA release significantly.

**Figure 1.**
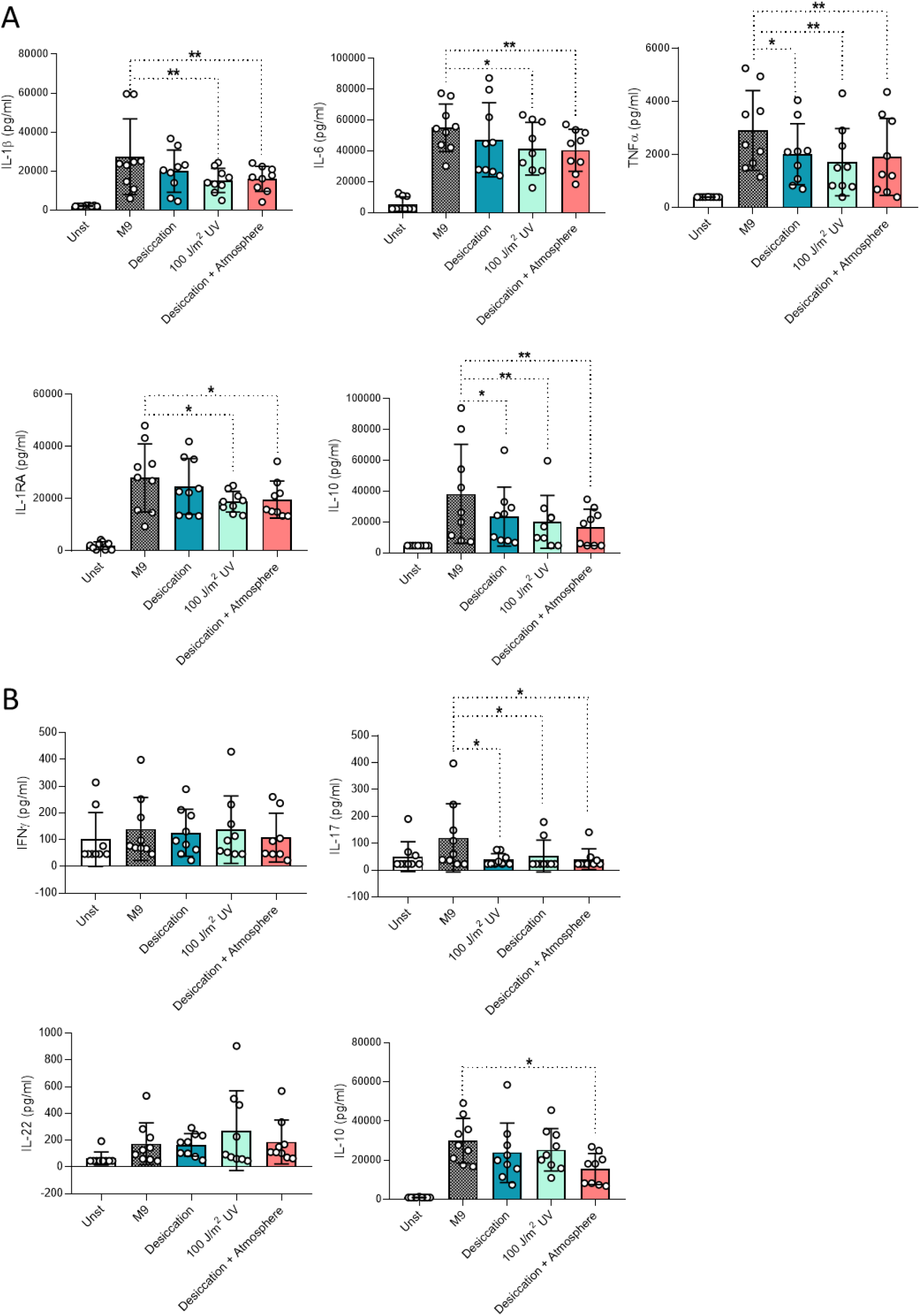
Cytokine production of PBMCs incubated with *K. pneumoniae* previously exposed to Mars-like conditions. PBMC innate (A) and adaptive (B) cytokine responses. n=9 biological replicates pooled from 3 independent experiments, *p≤0.05, **p≤0.01, Friedman test followed by Dunn’s multiple comparison test. Unst: unstimulated, M9: minimal media, UV: ultra-violet.

Cytokine release in response to *S. marcescens* exposed to simulated Mars conditions differed from *K. pneumoniae*, with reduced IL-1β and TNFα upon desiccation alone or combined with Martian atmosphere or UV radiation (Figure 2A). For all exposure regiments, the release of IL-6 and IL-10 did not show differences compared to the unexposed controls. Notably, unlike *K. pneumoniae*, IL-1RA release increased significantly when *S. marcescens* was exposed to desiccation. The production of T-cell-derived cytokines IFNγ and IL-10 was lower upon stimulation with *S. marcescens* exposed to desiccation alone and desiccation combined with UV radiation when compared with the standard-grown microorganism, while IL-22 secretion was not altered and IL-17 production could not be detected (Figure 2B). Exposures of *S. marcescens* to UV radiation (400 J/m^2^) did not alter cytokine induction in human immune cells, compared to control bacteria.

**Figure 2.**
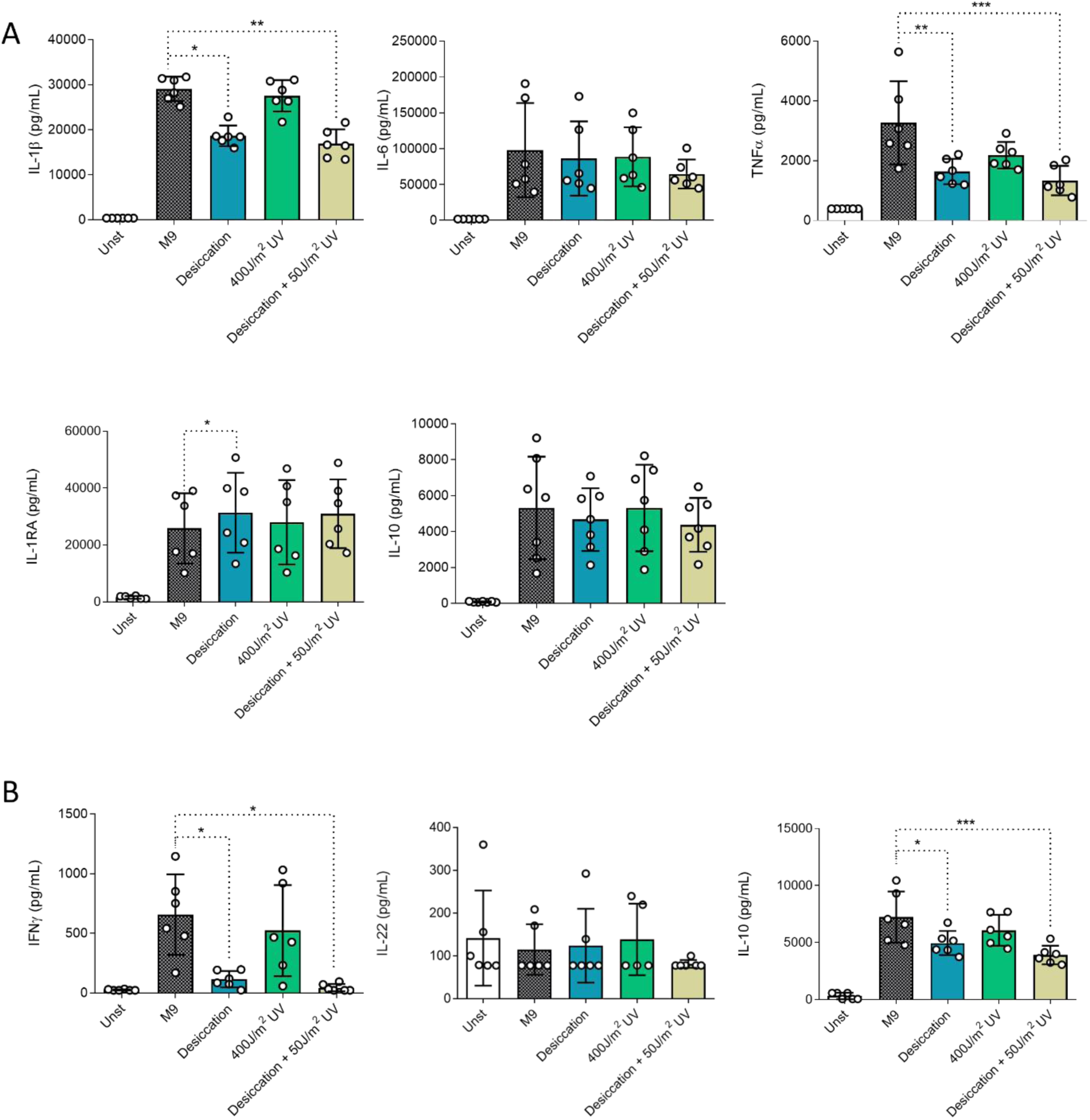
Cytokine production of PBMCs incubated with *S. marcescens* previously exposed to Mars-like conditions. PBMC innate (A) and adaptive (B) cytokine responses. n=6 biological replicates from 2 independent experiments, *p≤0.05, **p≤0.01, ***p≤0.001, Friedman test followed by Dunn’s multiple comparison test. Unst: unstimulated, M9: minimal media, UV: ultra-violet.

Taken together, our results show that the cytokine release profile triggered by the bacterial species exposed to Mars-like conditions differ between the two species tested. However, the exposure to the desiccation condition altered the cytokine response (TNFα, IL-10, IL-17 for *K. pneumoniae* and IL-1β, TNFα, IL-1RA, IFNγ and IL-10 for *S. marcescens*) for both bacterial species. In contrast, the effects of exposure to polychromatic UV radiation were not shared, with a reduction in the cytokine response (IL-1β, IL-6, TNFα, IL-1RA, IL-10 and IL-17) only seen for *K. pneumoniae*, despite the lower dose of UV exposure.

### Bacterial species exposed to Mars-like conditions showed altered capacity to induce reactive oxygen species and phagocytosis

To more broadly understand the innate immune responses to simulated Mars condition-exposed *K. pneumoniae* and *S. marcescens*, we quantified the production of ROS by PBMCs incubated with the bacteria (Figure 3A and B). Desiccation reduced ROS production induced by both species compared to standard-grown microorganisms, while UV exposure showed no effect. Similarly, a significant reduction in ROS production by PBMCS was seen for *K. pneumoniae* exposed to desiccation and pressure in Mars atmosphere and in *S. marcescens* exposed to desiccation coupled to UV radiation. These reductions in ROS and cytokine production indicate impaired immune recognition of Mars-like condition adapted bacterial species. To further elucidate innate responses upon a potential infection on Mars, the phagocytic capacity of CD11b+ cells was evaluated. All the exposure conditions tested altered phagocytic capacity compared to the control for both bacterial species. All simulated Mars conditions led to enhanced *K. pneumoniae* phagocytosis by human CD11b+ immune cells (Figure 3C and D). In contrast, the phagocytosis of *S. marcescens* was significantly reduced upon exposure to desiccation or to UV radiation (Figure 3E and F). These differences in phagocytic capacity indicate that *K. pneumoniae* and *S. marcescens* adapt distinctively to Mars-like conditions, suggesting divergent phenotypic changes between the two species.

**Figure 3.**
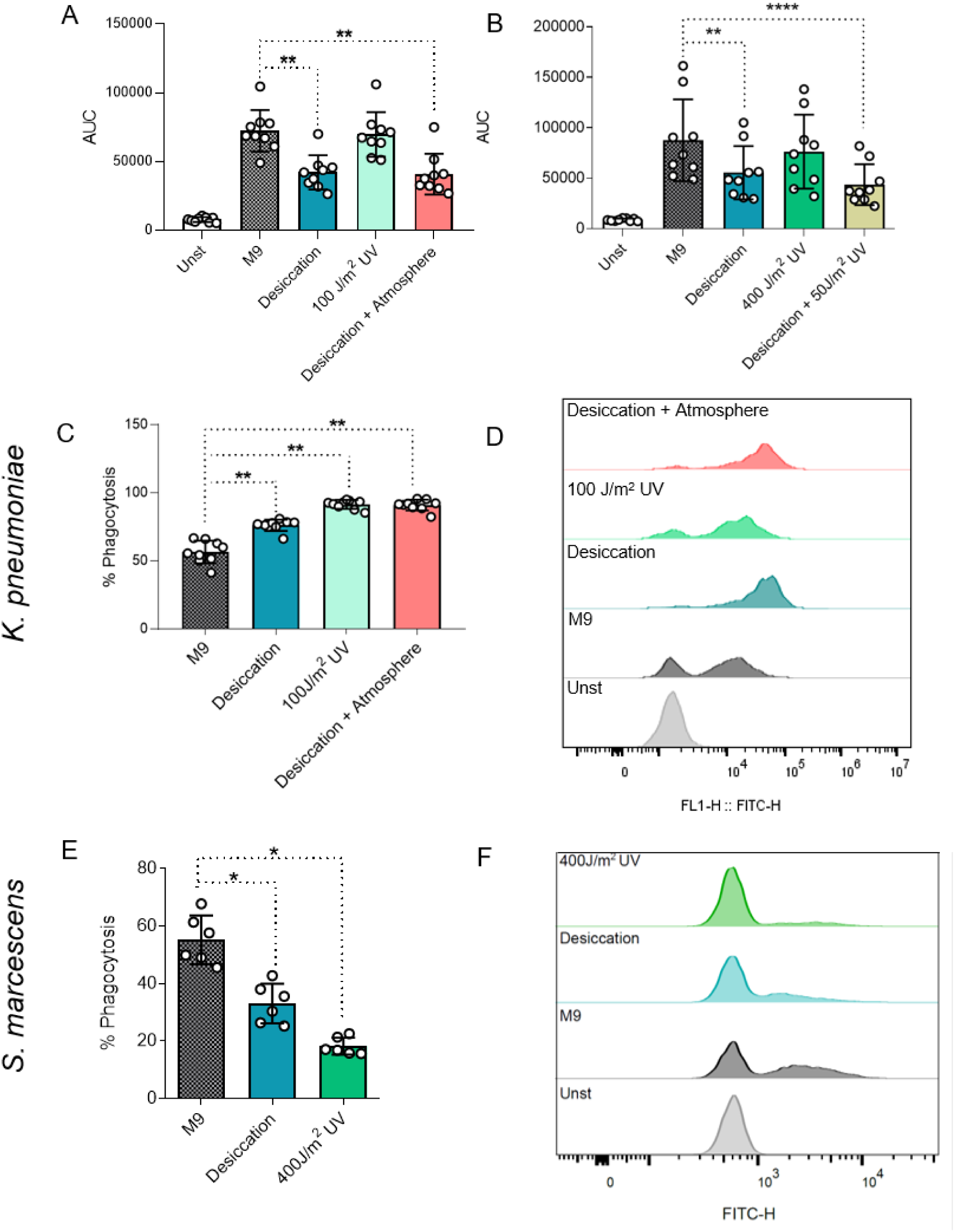
Effector functions of PBMCs against bacteria exposed to Mars-like conditions. Reactive oxygen species (ROS) production by PBMCs stimulated with *K. pneumoniae* (A) or *S. marcescens* (B). Percentage of CD11b+ cells positive for DTAF (FITC)-labelled bacteria (C and E) and representative histograms (D and F) for *K. pneumoniae* and *S. marcescens*. n=6-9 biological replicates from 2-3 independent experiments, *p≤0.05, **p≤0.01, ****p≤0.0001 Friedman test followed by Dunn’s multiple comparison test. Unst: unstimulated, M9: minimal media, AUC: area under curve as a ratio of ROS production over time, UV: ultra-violet.

### Desiccation-induced changes are partially reversible upon bacterial regrowth

Bacterial species with pathogenic potential exposed to Mars-like conditions may pose a health risk not only during human visitation to the planet, but also upon return to Earth. Therefore, we evaluated whether desiccated bacteria retain their altered phenotype after regrowth in minimal media. Regrowth of desiccated *K. pneumoniae* restored IL-10 production, while TNFα release remained unchanged (Figure 4A). Notably, exposure to desiccated *K. pneumoniae* did not significantly reduce TNFα secretion in this set of experiments in contrast to Figure 1. Regrowth also reversed ROS reduction and restored phagocytosis of *K. pneumoniae* (Figure 4C and D). Similarly, regrowth of *S. marcescens* restored IL-1β and IFNγ production to control levels (Figure 4E and F). However, TNFα and IL-1RA production stimulated by *S. marcescens* remained altered after regrowth, indicating partial reversibility of the immune response upon regrowth. *S. marcescens* also showed recovery in ROS production and phagocytosis upon regrowth (Figure 4G and H). Taken together, these results suggest that desiccation induces partially reversible phenotypic changes in both *K. pneumoniae* and *S. marcescens*, and highlight the bacteria’s ability to rapidly adapt after exposure.

**Figure 4.**
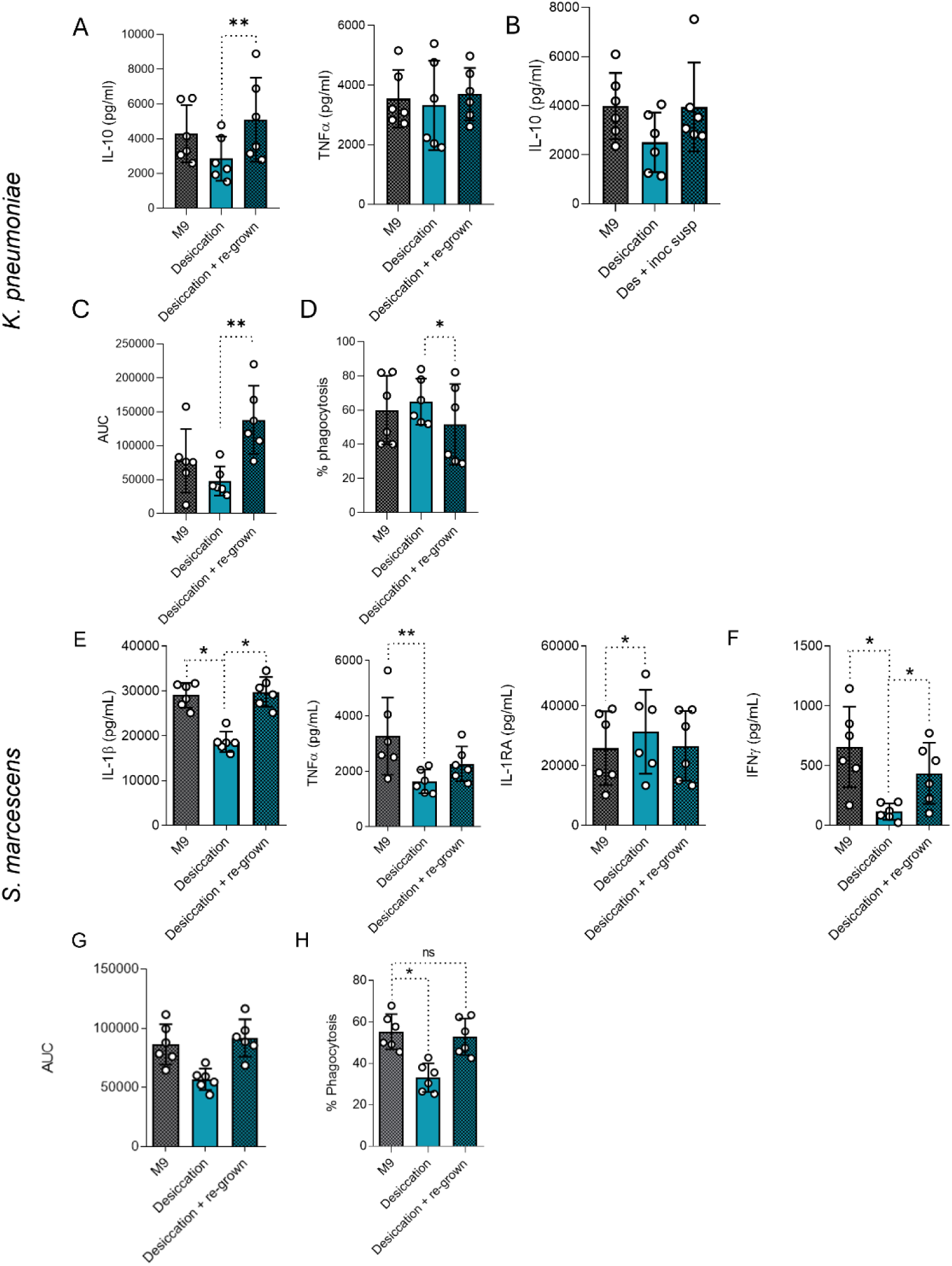
Re-growth after desiccation partially rescues the effects of desiccation. Cytokine production following stimulation of PBMCs for 24 hours (A, E) and 7 days (B, F) with *K. pneumoniae* (A, B) or *S. marcescens* (E, F). Reactive oxygen species production when stimulated with *K. pneumoniae* (C) or *S. marcescens* (G). Phagocytosis of *K. pneumoniae* (D) and *S. marcescens* (H) by CD11b+ cells n=6 biological replicates from 2 independent experiments. *p≤0.05, **p≤0.01, Friedman test with Dunn’s multiple comparison test, M9: minimal media.

### Desiccation reduced bacterial size and complexity

Morphological changes in bacteria exposed to Mars conditions likely influence immune responses. We investigated the cell morphology of *K. pneumoniae* and *S. marcescens* after exposure to desiccation, UV radiation, desiccation and Mars atmosphere and desiccation and regrowth, using transmission electron microscopy (TEM) (Figure 5A and C). TEM revealed distinct morphological changes in both bacterial species indicating cell envelope stress as well as a large population of cells with reduced electron density. *K. pneumoniae* cells remained rod-shaped post-desiccation, but had an irregular and shriveled appearance after UV exposure or combined conditions (desiccation and atmosphere and desiccation and regrowth). In contrast, *S. marcescens* generally retained its shape, showing irregularities resembling membrane blebbing only after desiccation and regrowth.

**Figure 5.**
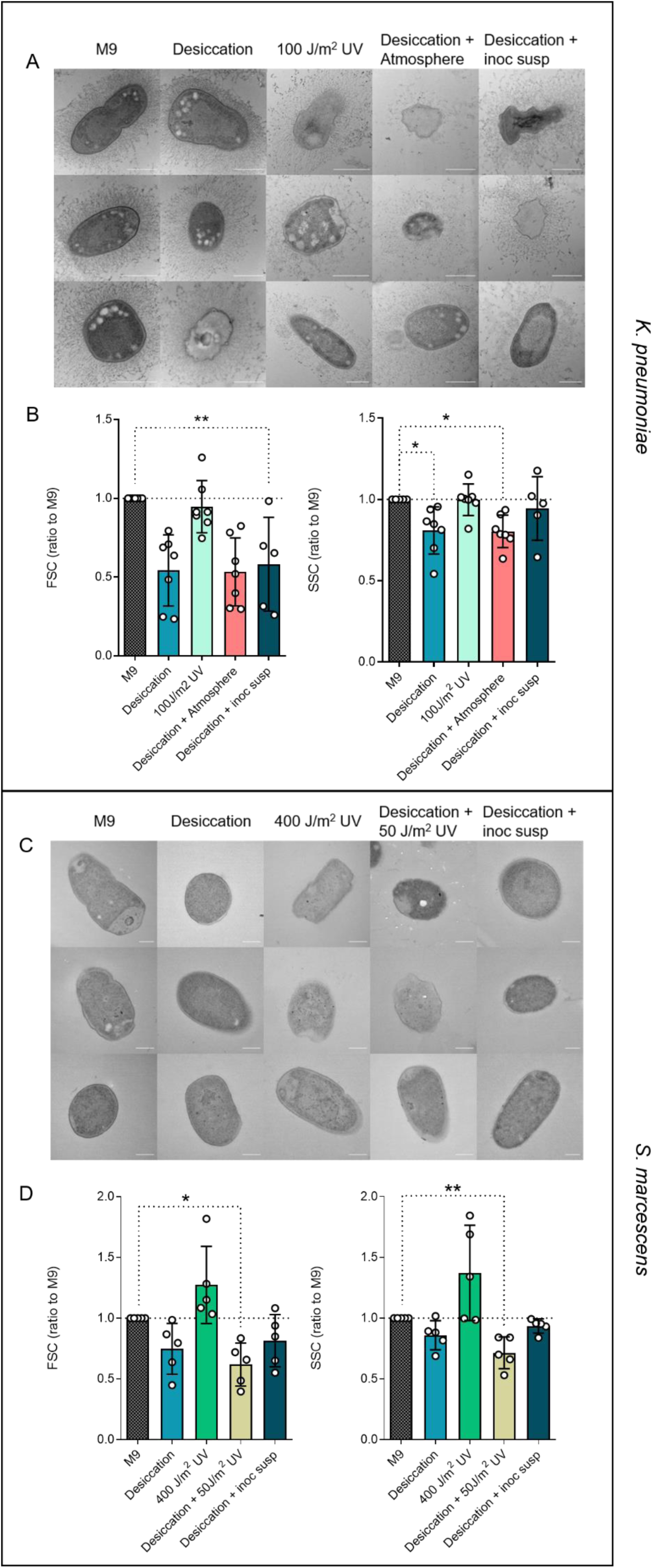
The morphology of bacterial cells is altered following exposure to simulated Mars conditions. TEM visualization of *K. pneumoniae* (A) and *S. marcescens* (C), size (FSC) and complexity (SSC) quantification by flow cytometry of *K. pneumoniae* (B) and *S. marcescens* (D). Scale bars are 500 nm in panel A and 250 nm in panel C. n=5-7 biological replicates from 2 independent experiments. *p≤0.05, **p≤0.01 Friedman test followed by Dunn’s multiple comparison test, M9: minimal media, UV: ultra-violet.

To further characterize bacterial morphology, we quantified the bacterial size (forward scatter, FSC) and complexity or granularity (side scatter, SSC) by flow cytometry. UV radiation had no impact on size or complexity, but both bacterial species showed size and complexity variation after desiccation alone or combined with other Mars-like conditions. Notably, the reduced size and complexity of desiccated *K. pneumoniae* and *S. marcescens* persisted after regrowth. These findings suggest that desiccation induces persistent morphological changes which might affect the recognition and response by immune cells.

## Discussion

Understanding how Earth-born pathogens exposed to Mars conditions interact with the human immune system is vital for assessing risks to crewed Mars missions. Our findings show that Mars-like conditions alter the morphology of the *K. pneumoniae* and *S. marcescens* cells and subsequently the interaction with immune cells, affecting cytokine release, phagocytosis, and ROS production, all important antimicrobial mechanisms. The immune response differences are likely driven by the specific stressors applied to the bacterial cells. Cell inactivation by Mars-like stressors, as shown in our previous research (Zaccaria et al., 2024), may also impact cytokine responses. Research in LEO has demonstrated changes in bacterial growth rates, drug interactions, and cell structure (Higginson et al., 2016, Sharma and Curtis, 2022), similar to the alterations observed in our study. Our findings provide a valuable foundation for understanding the potential infection risks during crewed Mars missions, emphasizing the need for further investigation.

Mars conditions may alter molecules on the surface of bacteria and the structure of the bacterial cell walls and other cell surface structures, impacting their interactions with immune cells (Wilson et al., 1998). Epithelial and immune cells express pattern recognition receptors (PRRs) that, upon detecting foreign material, trigger cytokine release and further activation of the immune system. Variations in cytokine and chemokine responses can lead to the altered activation of immune cells (Muri et al., 2023). Human immune cells are also affected by space conditions (e.g. low gravity), however this interaction was mostly studied in the context of the adaptive immune system (Akiyama et al., 2020). Concerning the innate immune cells, we know that the number of neutrophils increased at landing compared to pre-flight levels, while phagocytosis and oxidative burst capacities were significantly lower than control mean values after 9- to 11-day missions (Kaur et al., 2004). Also changes in monocytes have been studied in astronauts participating in spaceflights, showing that monocytes exhibited reductions in the ability to engulf *Escherichia coli*, to elicit an oxidative burst, and to degranulate (Kaur et al., 2005). Furthermore, when challenged with endotoxin (LPS), the space flight crew member’s monocytes collected at different time points produced lower amounts of interleukin-6 (IL-6) and IL-1β and higher levels of IL-1RA and IL-8 compared to those of control subjects (Kaur et al., 2008). Immune dysregulation during spaceflight, combined with microbial presence in spacecraft (Fernandez-Gonzalo et al., 2017, Checinska Sielaff et al., 2019, Capri et al., 2023, Jacob et al., 2023) and microbiome changes (Tierney et al., 2024), may pose significant health risks and treatment challenges. Earlier studies have shown that during *K. pneumoniae* lung infection the concentrations of IL-1β, IL-6 and TNFα are induced, while TNFα neutralization increased mice mortality (Wei et al., 2023). Therefore, reductions in cytokine release observed may indicate bacterial evasion strategies to escape immune detection. This suggests that the exposure to the different environmental Mars conditions might alter bacterial pathogenicity.

The altered cytokine and ROS profiles indicate that the immune system may have impaired the ability to recognize Mars-adapted bacteria. Defective cytokine stimulation could also decrease other immune processes, including phagocytosis and bacterial killing (Winterbourn et al., 2016, Canton et al., 2021). To assess whether these changes induced by the Mars-like conditions would persist, we assessed PBMC reactions to desiccated and subsequently regrown bacterial cells. Our findings show that regrowth of desiccated bacteria partially restores the immune system’s ability to recognize them. Further studies are needed to assess immune responses to bacteria regrown after exposure to other Mars-like conditions or multiple desiccation cycles, providing valuable insights for long-term Mars missions.

Although Mars conditions were simulated in this study, they also provide valuable insights applicable to Earth-based scenarios. Desiccation, known to cause significant cellular damage, is particularly relevant. Many microbial species exhibit desiccation survival, which facilitates pathogen transmission (Salazar et al., 2022). This survival adaptation enables pathogens to spread and increase host infection risks (Green et al., 2022, Javed et al., 2024). Thus, desiccation is not only significant for astrobiology, but also for public health, particularly in understanding bacterial spread in hospitals and public transport. Despite its importance, studies on immune recognition of desiccated bacteria remain limited. Our study demonstrated that desiccation decreased bacterial size and complexity, aligning with previous findings that oxygen deprivation reduces cell size (Bashiri et al., 2021). Under Mars-like low-pressure conditions, further size reduction in *K. pneumoniae* was not observed, likely due to the already strong effect of desiccation. Transmission electron microscopy revealed heterogeneity among treatment groups. Understanding these morphological changes is crucial for developing sterilization techniques for both hospitals and spacecrafts (Spry et al., 2024).

This study revealed that Mars adaptation of bacteria significantly alter immune responses, offering fundamental insights into potential health risks for future crewed Mars missions. While our findings are derived from healthy donor samples on Earth, evaluating immune reactions of immune cells isolated from astronauts during or after spaceflight would provide a more precise understanding of the pathogenic threats posed by these adapted microorganisms. These results highlight the urgent need for proactive risk prevention strategies, both for crew safety and planetary protection. Comprehensive immunological measures are essential throughout all mission phases: departure, travel and habitation on Mars. Implementing rigorous protocols to control bacterial spread on both internal and external surfaces of habitats must be a top priority. As we prepare for the complexities of long-duration space missions, early identification and mitigation of these health risks are crucial to ensure mission success and the well-being of both crew members and the humans who will come into contact with them upon their return to Earth.

## Materials and Methods

### Bacterial strains and growth media

The bacterial species *K. pneumoniae* 298-53 (DSM 30104) and *S. marcescens* BS303 (DSM 30121) were purchased from Leibniz Institute DSMZ-German Collection of Microorganisms and Cell Cultures GmbH. Following growth in Nutrient Broth (Difco; NB) and on NB agar (15 g/L), at 35°C overnight, colonies from each species were incubated in M9-complete minimal salts media (47.8 mM Na_2_HPO_4_, 22.0 mM KH_2_PO_4_, 8.6 mM NaCl, 3.7 mM NH_4_Cl, 2 mM MgSO_4_, 0.1 mM CaCl_2_ and 0.02 mM Fe(III)Cl_3_ without carbon source) as described previously by Domínguez-Andrés et al. (2020). Growth curves were plotted by measuring the optical density at 600 nm (OD_600_) of the bacterial culture following the dilution of 0.2% (w/v) D-gluconic acid sodium salt (C_6_H_11_NaO_7_), (Fluorochem), and D-glucose monohydrate (C_6_H_12_O_6_), (Merck), in the M9-complete media. Liquid NB and M9-complete media with D-glucose were used as positive controls, instead M9-complete without carbon source was used as negative control. The Mars-like exposure regimens were selected based on Mars simulated conditions which affected the survival of the bacteria while retaining viability of the cultures as described in Zaccaria et al. (2024).

### Exposure to desiccation

To understand the effect of desiccation on the immune response to the bacteria, late exponential phase cultures of *K. pneumoniae* and *S. marcescens* were grown in M9-gluconic medium. For each species, aliquots of 200 µl were placed on sterile 1 cm diameter glass disks and left to dry in a running clean bench at room temperature for 24h, under sterile conditions with a relative humidity of ca. 30% ± 10%. For processing, the cells were resuspended in 5 ml Eppendorf tubes containing 1 ml of PBS, lightly vortexed and allowed to soak for 1h at room temperature.

### Exposure to polychromatic UV radiation

The effect of UV radiation was evaluated by growing *K. pneumoniae* and *S. marcescens* in M9-gluconic to late exponential phase. In closed sterile UV transmissible quartz cuvettes (0.5 cm path-length, Hellma GmbH & Co. KG, Muelheim, Germany), a volume of 3 ml of each bacterial culture was placed. The cuvettes contained a magnetic stirrer allowing all the cells to be stirred while being irradiated. The cuvettes were placed vertically at 112 cm from the light source. Each sample was irradiated with polychromatic UV (200-400 nm) with fluences of 10.35 W/m^2^ using the SOL2 polychromatic UV lamp, equipped with a UV 500S irradiation source (Dr. Hoenle AG, UV-Technologies, Germany) at the German Aerospace Center in Cologne, Germany. The fluence of the lamp was determined with the Bentham DMc150 transportable spectroradiometer (Bentham Instruments Ltd., Reading, UK), with optics inside the simulation chamber as per Rabbow et al. (2005). Following irradiation liquid cultures were visually evaluated under a light microscope (Zeiss Primostar, Carl Zeiss Microscopy GmbH, Jena, Germany) to identify a potentially altered morphology.

### Exposure to Mars atmosphere and pressure

The species were exposed to Mars-like atmosphere composed of a Mars-like gas mix (2.7% N_2_, 1.6% Ar, 0.15% O_2_ in CO_2_ vol/vol) and Mars-relevant pressure of 6 hPa using a vacuum pump (Rotary vane pump DUO 035 and HIPace 700, Pfeiffer Vacuum GmbH, Germany) monitored constantly during the exposure (TPG 262 Full Range Gauge, Pfeiffer Vacuum GmbH, Germany). The samples were exposed to Mars atmosphere following desiccation as described in 2.2. The desiccated glass disks were placed in the gastight Trex-box (Transport and exposure-box, Beblo-Vranesevic et al. (2017)) and the Mars atmosphere at Mars-relevant pressure applied.

The bacterial species were exposed to the described conditions in order to replicate Mars-relevant conditions and at the same time to have enough cell survival to potentially determine infection of the human body and to assess the immune response.

### Inactivation of the bacteria

Immunological evaluations were performed at Radboudumc, Nijmegen, the Netherlands. The bacteria were shipped inactivated using β-propiolactone (BPL) as follows. A citrate buffer was prepared with 125 mM sodium citrate (C_6_H_5_Na_3_O_7_ x 2H_2_O; Sigma-Aldrich) and 150 mM sodium chloride (NaCl, Merck). A 10% (v/v) stock solution of 98% BPL (Acros Organics), in citrate buffer was prepared. The 10% stock solution was added to the exposed bacteria to a final concentration of 0.1% (v/v) of BPL. The bacterial samples were stored at 4°C until processing at Radboud UMC. The samples were then incubated at 37°C for 4 hours to inactivate the BPL.

### Isolation of human peripheral blood mononuclear cells (PBMCs)

Buffy coats from healthy donors were obtained after written informed consent (Sanquin Blood Bank, Nijmegen, Netherlands). Samples were anonymized to safeguard donor privacy. PBMC isolation was performed by differential density centrifugation over Ficoll-Paque (GE Healthcare). Cells were resuspended in RPMI 1640 medium (Dutch modified Invitrogen) supplemented with 5 µg/mL gentamicin (Centrafarm), 2 mM Glutamax (Gibco), and 1 mM pyruvate (Gibco).

### PBMC culture stimulation

0.5 x 10^6^ PBMCs per well were seeded in round bottom 96 well plates and stimulated with 10^6^ CFU/ml *K. pneumoniae* or *S. marcescens* for 24 hours or 7 days in RPMI medium supplemented with 10% human pooled serum. All conditions were tested for cytotoxicity by measuring lactate dehydrogenase in the conditioned media after 24h incubation according to the manufacturer’s instructions (CytoTox96, Promega).

### Cytokine quantification

Secreted cytokines were determined using commercial ELISA kits for IL-1β, IL-6, TNFα, IL-10, IL-1Ra, IL-17, IL-22 and IFNy (R&D Systems) following the instructions of the manufacturer. Concentrations lower than the detection limit of the ELISA were replaced with the lowest detectable concentration for statistical analyses.

### Reactive oxygen species (ROS) quantification

Superoxide anion levels were evaluated using luminol-enhanced chemiluminescence and determined in a luminometer (Biotek Synergy HT). A total of 0.5 x 10^6^ PBMCs per well were incubated with serum opsonized 10^6^ CFU/ml *K. pneumoniae* or 10^7^ CFU/ml *S. marcescens*. Luminol (Sigma) was added to each well to start the chemiluminescence reaction. Each measurement was carried out in quadruplicates. Chemiluminescence was determined every 145 seconds at 37°C for 1 hour. Luminescence was expressed as relative light units (RLU) per second and as area under the curve (AUC).

### Phagocytosis assay

Bacteria were stained overnight with 100 µM Dichlorotriazinyl Aminofluorescein (DTAF) (ThermoFisher). After washing with PBS three times, stained bacteria were opsonized for 1 hour at 37°C in RPMI with 10% pooled human serum. 0.5 x 10^6^ PBMCs per well were plated in round bottom 96-well plates and incubated with *K. pneumoniae* or *S. marcescens* (MOI 2:1) for 15 minutes. Prior to flow cytometry analysis, cells were stained with anti-CD11b-BV785 (Biolegend) for 20 minutes at 4°C. Measurements were done using CytoFLEX (Beckman Coulter). Data presented as percentage of DTAF positive events within the CD11b+ population. Analysis was performed using FlowJo v10.9.

### Transmission electron microscopy (TEM)

*K. pneumoniae* and *S. marcescens* were grown overnight at 35°C in M9-gluconic medium and exposed to desiccation, polychromatic UV radiation or Mars atmosphere and pressure as described previously. The cell number of the bacteria for TEM was determined by CFU plating on NB agar (8 g/L). The cells were then fixed in a 4% solution of PFA (paraformaldehyde) in 0.1 M PHEM (PIPES, HEPES, EGTA and Magnesium Sulfate) buffer. An equal volume of PFA in PHEM buffer solution was added to unwashed cells following exposure to the listed conditions. The cells were allowed to fix at room temperature for 15 minutes with gentle agitation. The fixed cells were pelleted at 1500 G and the fixative was removed and replaced with fresh fixative. The cells were resuspended and allowed to fix for 40 minutes at room temperature with gentle agitation. The cells were then washed three times with PBS and stored in 0.1% PFA in PHEM at 4°C until being processed for TEM.

Pre-fixed cells were gently pelleted (600-RCF 5 min), after which supernatant was removed and pellet was resuspended in the remaining medium. Next, samples with a thickness of 100 µm were high-pressure frozen using the 3 mm platelet system in a HPM100 high-pressure freezer (Leica Microsystems). Samples were freeze-substituted in anhydrous acetone containing 2% OsO_4_, 1% H_2_O and 0.2% Uranyl acetate and subsequently embedded in Epoxy resin (van Niftrik et al., 2008). 60 nm sections were prepared on a Reichert-Jung Ultracut ultramicrotome and collected on 100 mesh copper grids with a carbon-coated formvar support film. Grids were post-stained with 0.5% Uranyl acetate and Reynolds lead citrate before analysis in the JEOL-1400 Flash TEM operating at 120 kV.

### Bacterial size and complexity quantification by flow cytometry

Bacteria were stained with Syto40 (ThermoFisher) to stain nucleic acids. Size and complexity are presented as geometric mean of FSC and SSC respectively, as a ratio to M9 control bacteria of Syto40+ events. Measurements were done using CytoFLEX (Beckman Coulter) and analysis performed using FlowJo v10.9.

## Abbreviations used

BPL: β-propiolactone
CFU: colony-forming unit
FSC: forward scatter
ISS: international space station
LEO: low earth orbit
NB: nutrient broth
PBMC: peripheral blood mononuclear cell PBS phosphate-buffered saline
PFA: paraformaldehyde
PHEM: PIPES, HEPES, EGTA and Magnesium sulfate buffer PVP polyvinyl-pyrrolidone
ROS: reactive oxygen species
SSC: side scatter
TEM: transmission electron microscopy
UMC: university medical center
UV: ultraviolet

## Acknowledgements

We thank Andre Parpart for help and guidance in setting up the atmosphere exposure experiment, Hetty Manenschijn for maintenance of TEMs at the department of general instrumentation (Faculty of Science, Radboud University).

## Conflict of interest

No competing financial interests exist related to the work presented in this study. MGN is scientific founder of Biotrip, TTxG, Lemba and Salvina.

## Funding Information

This work was supported by an Off Road ZonMw Grant awarded to J.D.-A (04510012010022). M.G.N. was supported by an ERC Advanced Grant (833247) and a Spinoza Grant of the Netherlands Organization for Scientific Research.

